## Supplemental Figures_revised version for "On the Role of VP3-PI3P Interaction in Birnavirus Endosomal Membrane Targeting"

Laura Ruth Delgui

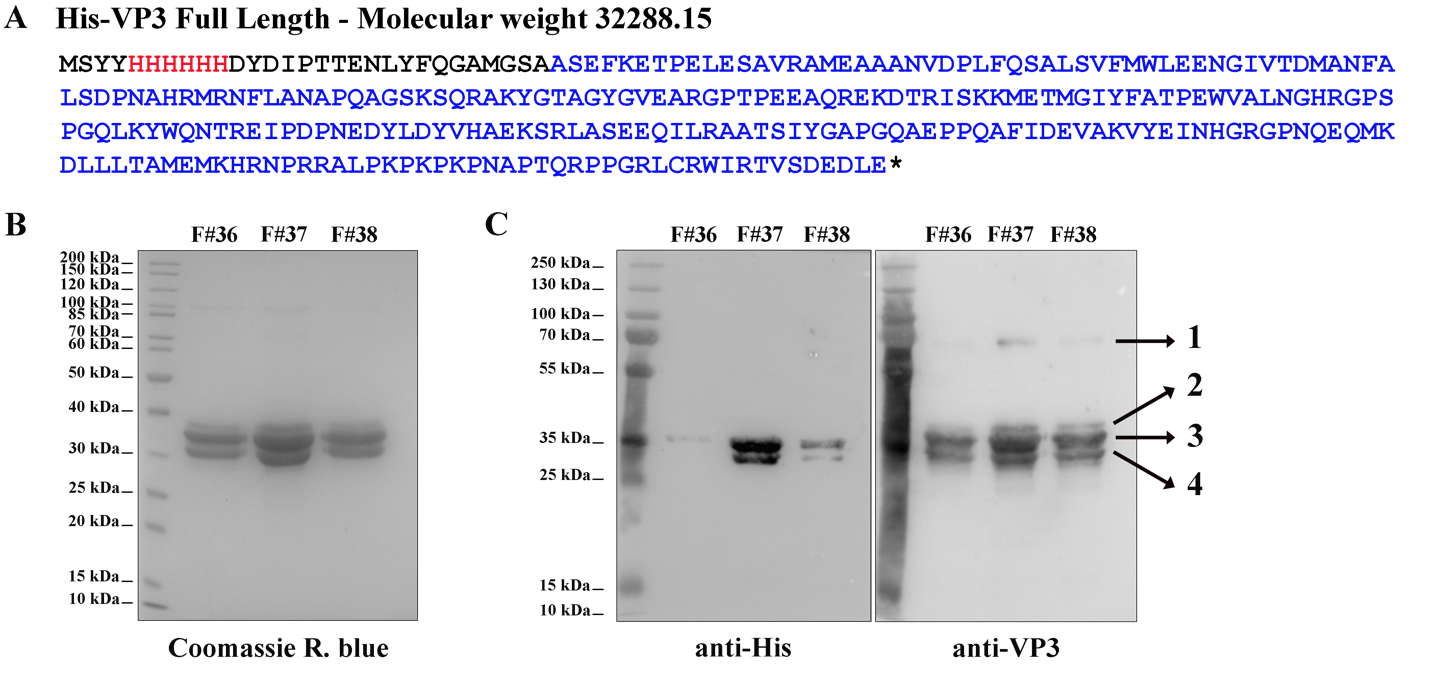

**Fig. S1. His-VP3 Full length (FL) purification and characterization.**

**(A)** The aminoacidic sequence of His-VP3 FL and its theoretical molecular weight. The His-tag is colored in red and the VP3 sequence in blue. Additional residues are black.

**(B)** Coomassie R. blue-stained polyacrylamide gel showing fractions (F) #36, #37 and #38 from the size exclusion chromatography.

**(C)** Western blot images (left: anti-His; right: anti-VP3) from gels identical to that shown in (B). Numbers 1 to 4 indicate the identified proteins by mass spectrometry. 1. His-VP3 FL dimer (~70 kDa); 2. Sulfonylated His-VP3 FL (due to the use of phenylmethylsulfonyl fluoride as proteases inhibitor); 3. His-VP3 FL; 4. His-VP3 lacking the last 14 Ct residues.

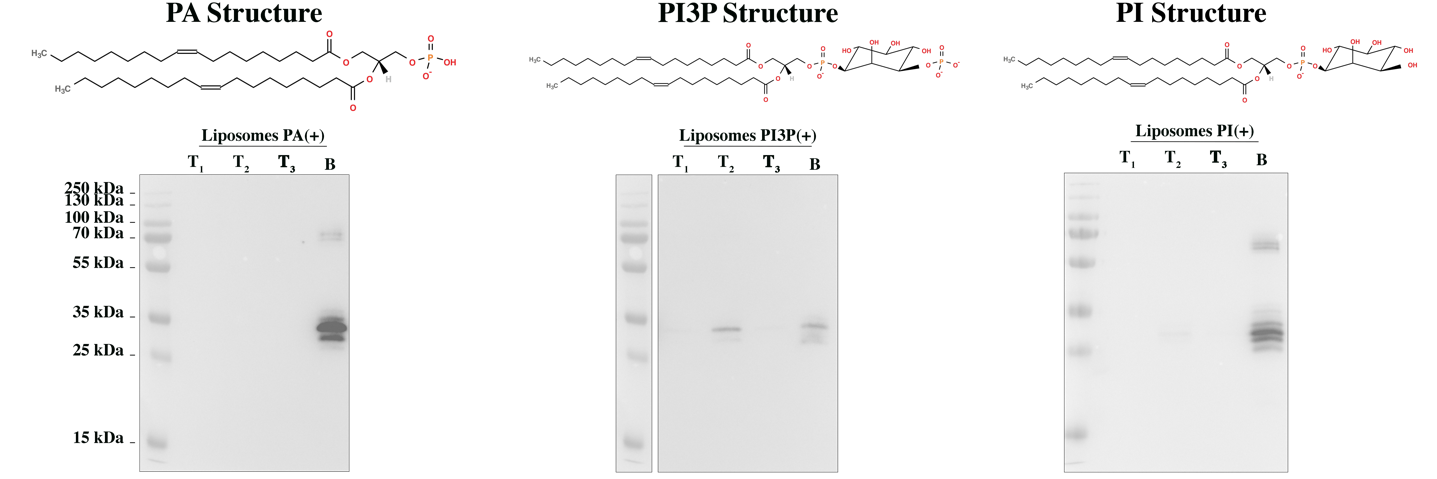

**Fig. S2. His-VP3 FL PI3P-binding specificity.**

Immunoblots of the three top (T_1_, T_2_ and T_3_) and bottom (B) fractions from co-flotation assays using liposomes where PI3P was replaced by 1,2-dioleoyl-sn-glycero-3-phosphate (PA) or [1,2-dioleoyl-sn-glycero-3-phospho-(1'-myo-inositol)] (PI) in an identical molar ratio, named “liposomes PA(+) or PI(+)”. We observed that while His-VP3 FL bound to liposomes PI3P(+) (middle panel), it did not bind to liposomes PA (left panel) or PI (right panel), reinforcing the notion that the interaction of VP3 with PI3P is specific. On top of each panel, the corresponding lipid structure is shown.

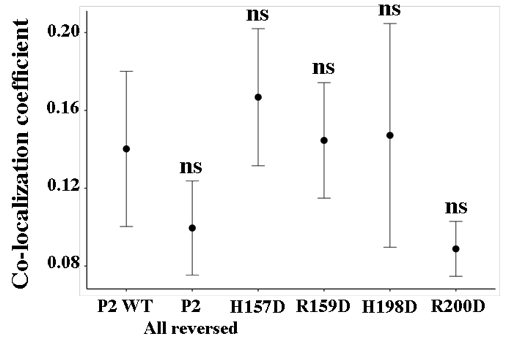

**Fig. S3. VP3-EGFP-Rab5 co-localization quantification.**

The dot plot depicts the co-localization coefficient for each protein determined as explained in the Materials and Methods section. Significant differences (ns *P* >0.05) as determined by one-way ANOVA with Tukey’s HSD test.

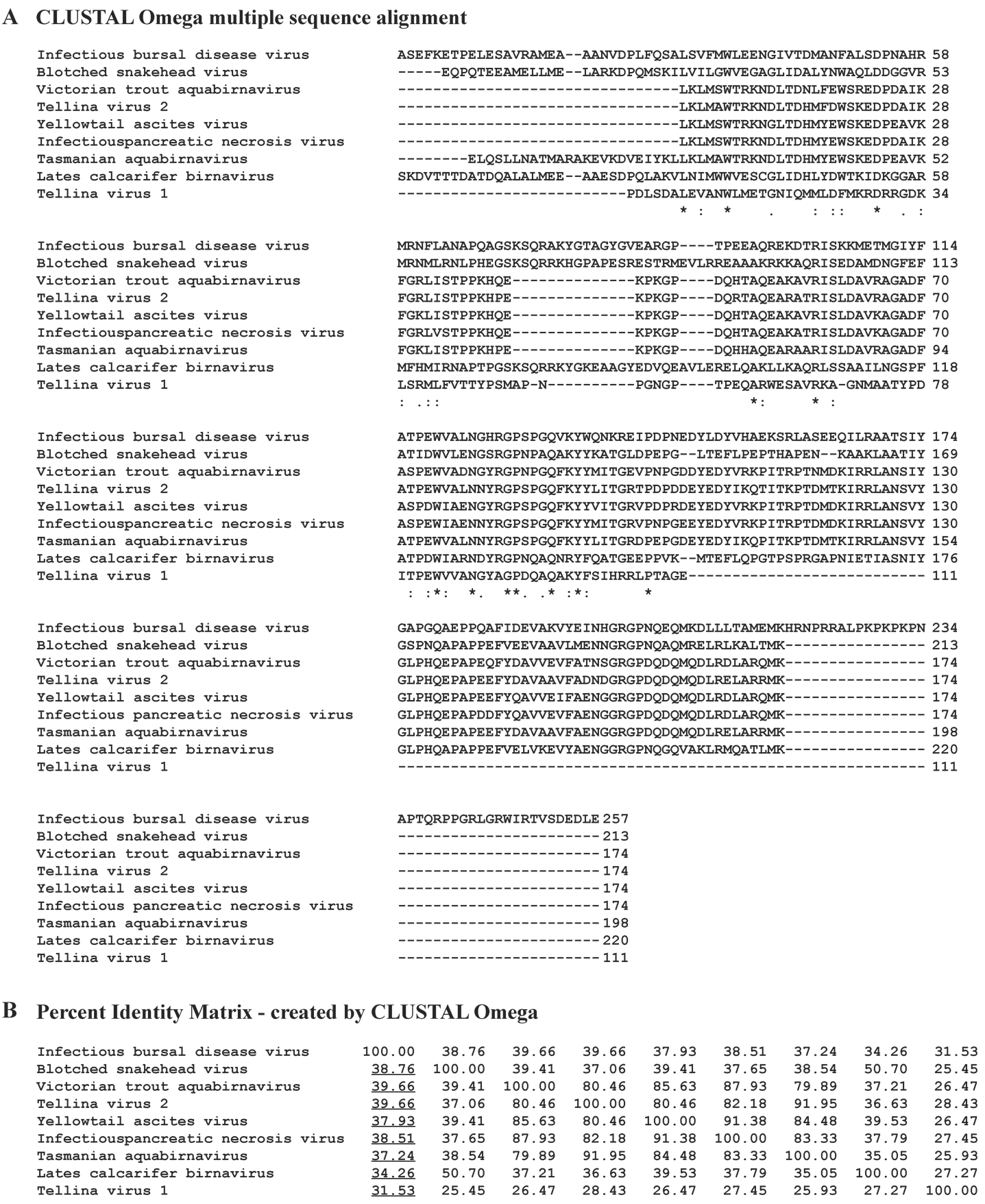

**Fig. S4. Bioinformatic analysis.**

**(A)** VP3 multiple sequence alignment analysis performed with CLUSTAL Omega (Sievers et al., 2011) with IBDV VP3 (AF_140705.1) as the reference. Characters "*", ":" and "." below the sequences indicate identity (100% of conservation), homology (strongly similar, > 50% of conservation) and homology (< 50% of conservation), respectively.

**(B)** Matrix showing the percentages of identity of non-IBDV VP3 related to IBDV VP3 (AF_140705.1) performed with CLUSTAL Omega as well. Underlined are indicated those of non-IBDV VP3 related to IBDV VP3.

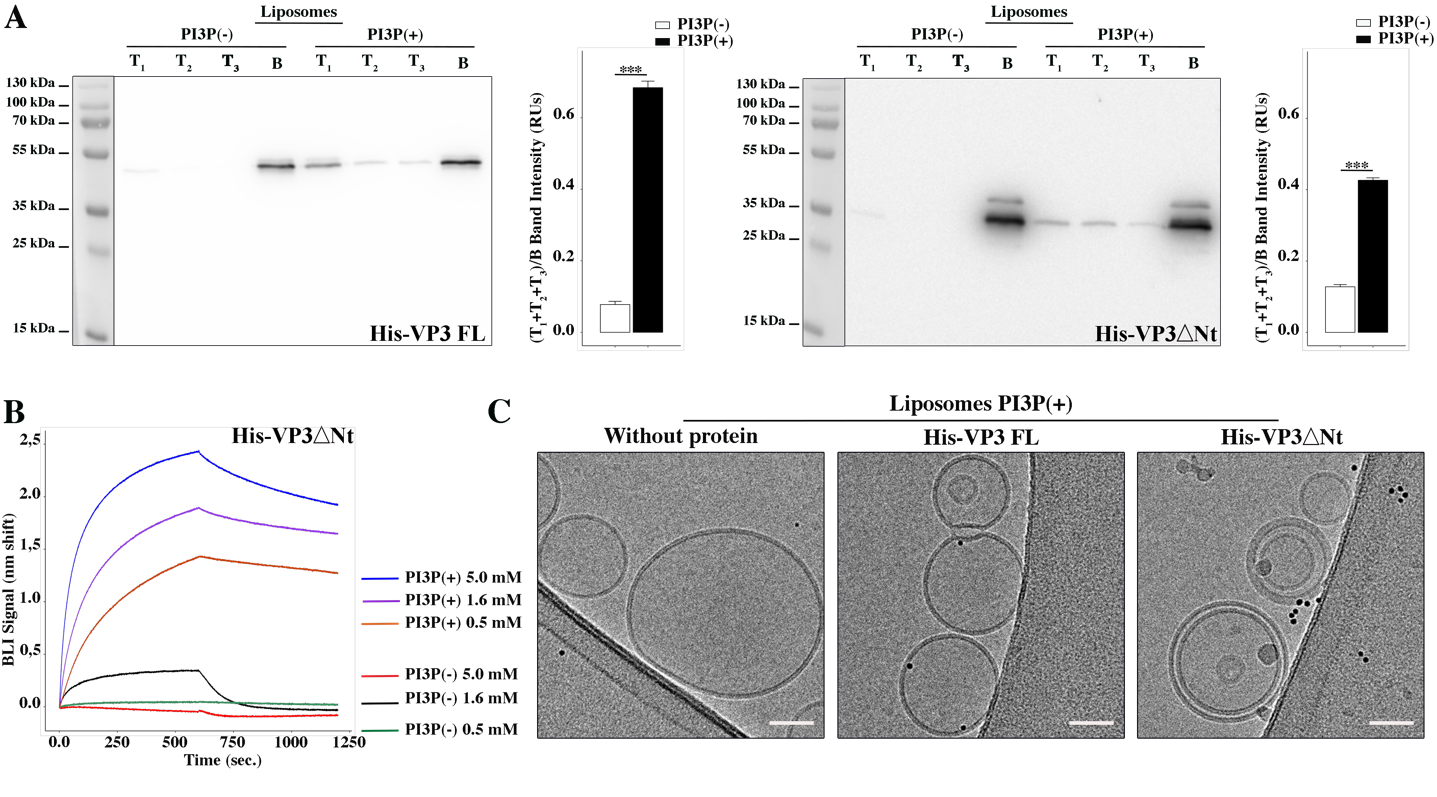

**Fig. S5. Biophysical characterization of His-VP3 ΔNt binding to PI3P.**

**(A)** Left panel: Immunoblots of the three top (T_1_, T_2_ and T_3_) and the bottom (B) fractions from a liposome PI3P(-) or PI3P(+) OptiprepTM co-floatation assay indicating that His-VP3 FL protein specifically binds to liposome PI3P(+). Results are representative of three independent experiments. The bar plot represents the intensity of (T_1_+T_2_+T_3_)/B bands for each liposome preparation. Right panel: Immunoblots of the three top (T_1_, T_2_ and T_3_) and bottom (B) fractions from a liposome PI3P(-) or PI3P(+) OptiprepTM co-floatation assay indicating that His-VP3 ΔNt protein specifically binds to liposome PI3P(+). Results are representative of three independent experiments. The bar plot represents the intensity of (T_1_+T_2_+T_2_)/B bands for each liposome preparation. Significant differences (*** *P* <0.001) as determined by one-way ANOVA with Tukey’s HSD test.

**(B)** Binding of His-VP3 ΔNt to three different concentrations of liposomes PI3P(-) or PI3P(+). Association and dissociation sensorgrams measured by BLI, showing the specific interaction of His-VP3ΔNt with liposomes PI3P(+) in a dose-dependent manner, as indicated.

**(C)** Transmission electron microscope images of cryo-fixated liposomes PI3P(+) control (without protein), or incubated with His-VP3 FL- or His-VP3 ΔNt-Ni-NTA gold particles showing electrodense particles decorating the membrane of the liposomes when His-VP3 FL or His-VP3 ΔNt were present. The scale bar represents 50 nm.

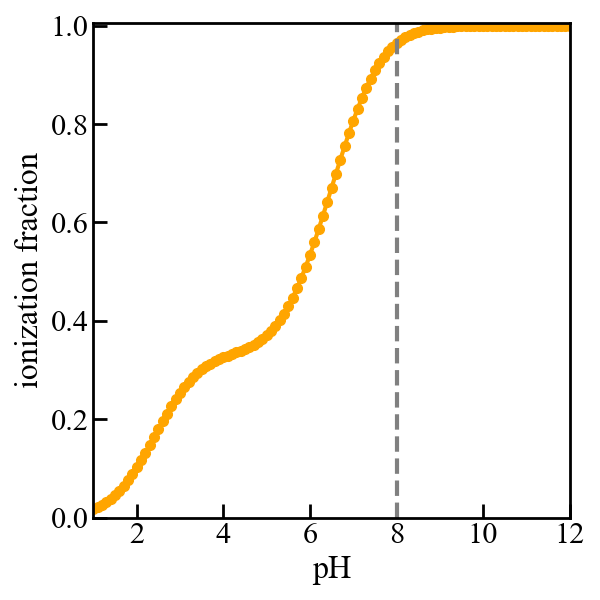

**Fig. S6. Titration curve of the membrane.**

Fraction of deprotonated (or ionized) acidic groups on the surface of the membrane versus pH of the bulk solution. The dashed vertical line indicates that at pH 8, the acidic groups are nearly fully deprotonated, resulting in a negative surface charge on the membrane. As mentioned in the manuscript, the model membrane consists of a 5 nm wide slab of dielectric material that exposes a mixture of 5% acidic and 95% neutral “head groups” to a 150 mM NaCl solution. The acidic lipids contain three acid moieties with pKa values of 2.5 and 6.5, as determined for PI3P. No proteins are present in the solution for the purpose of computing the titration curve. The curve was obtained using the Molecular Theory briefly described in the manuscript and fully detailed in the references (Chiarpotti et al., 2021; Ramírez et al., 2019).

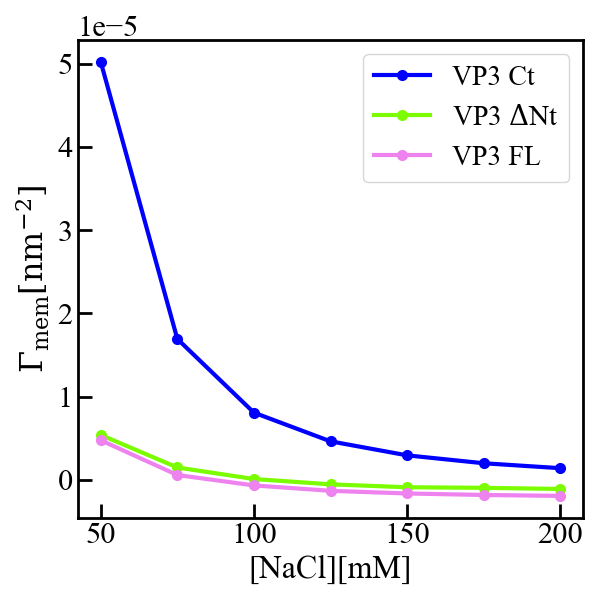

**Fig. S7. Interfacial concentration of VP3 constructs.**

Molecular Theory calculation of the surface concentration, or more precisely the surface excess concentration$\text{Γ}_{\text{mem}}$ of the three protein constructs VP3 Ct, VP3 ΔNt, and VP3 FL, as a function of the concentration of salt in the bulk solution. The overall electrical charge of each construct is +5, 0 and -3 respectively. The pH of the solution is 8, so that the surface of the membrane is negatively charged. The surface concentration of each protein decreases when increasing the concentration of salt, as the salt ions screen the electrostatic interactions between the proteins and the surface of the membrane. Clearly the effect is stronger for positively charged VP3 Ct. As detailed in reference (Chiarpotti et al., 2021) the surface excess is defined as $\text{Γ}_{\text{mem}}=\int_{0}^{\infty} \left[ \left\langle\rho_{p}(z) \right\rangle-\rho_{p}^{b} \right]dz$, where $\left\langle\rho_{p}(z) \right\rangle$ is the average protein density at position z, and $\rho_{p}^{b}$ is that in the bulk solution. $\text{Γ}_{\text{mem}}$ gives the total number of proteins per unit area, in excess of the bulk solution, that accumulate ($\text{Γ}_{\text{mem}}>0$ ) or deplete ($\text{Γ}_{\text{mem}}<0$) due the presence of the membrane.

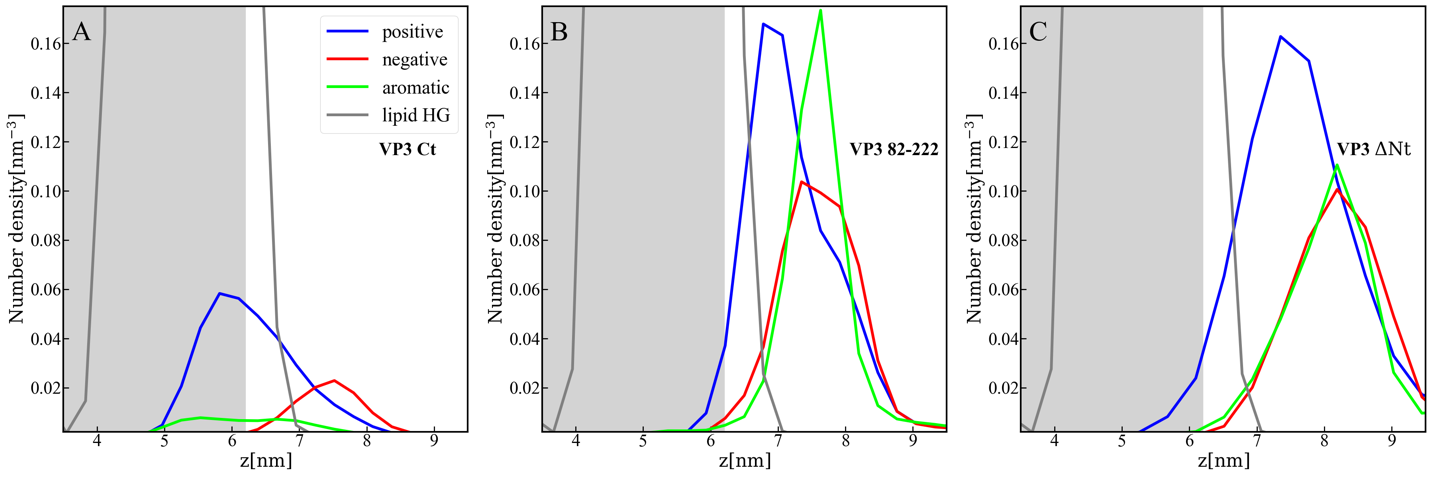

**Fig. S8. Distribution of charged and aromatic residues of VP3.**

Spatial distribution of charged groups obtained from coarse-grained Molecular Dynamics simulations of VP3 Ct (A), VP3 82-222 (B), and VP3 ΔNt (C) adsorbed on a lipid bilayer composed of DOPE, DOPC, and PI3P (with molar ratios of 64:31:5). DOPE and DOPC are electroneutral, while PI3P is anionic. VP3 82-222 represents a portion of the VP3 sequence for which a crystallographic structure is known (Gly82-Asn222). It was included in the plot for comparison with VP3 Ct (Arg223-Glu257) and VP3 ΔNt (Gly82-Glu257). The variable 'z' represents the z-component of the perpendicular distance between the center of mass of the membrane and the center of mass of each protein. The blue curves represent the distribution of positively charged residues in the protein, while the red curves represent the distribution of negatively charged residues. The green curves depict the distribution of aromatic residues, and the grey curves represent the distribution of the lipid's polar head groups. The grey shaded area schematically represents the region predominantly occupied by the lipids. As observed, in all cases, the positively charged amino-acids are in closer contact with the membrane surface and penetrate into the region of the lipid's polar head groups. In VP3 ΔNt, negatively charged and aromatic residues are somewhat more exposed towards the solution side of the interface.

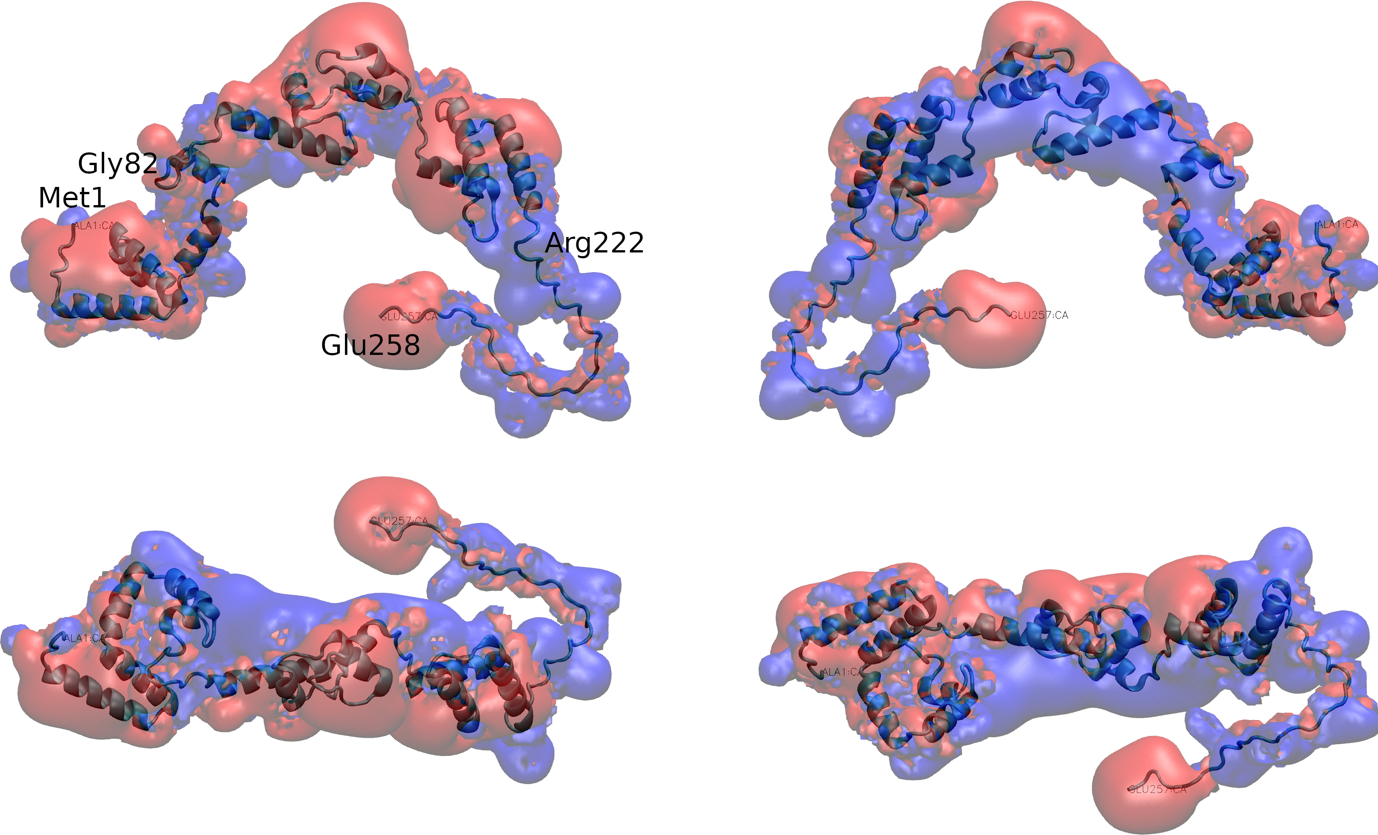

**Fig. S9. Map of electrostatic potential around VP3 FL.**

Four different views of the electrostatic potential map around VP3 FL, as computed with the Adaptive Poisson-Boltzmann Solver (APBS) plugin of VMD (Dolinsky et al., 2004). Electrostatic isopotential surfaces at +1.1 mV in red and at -1.1mV in blue account for negative and positive electrostatic potential, respectively.

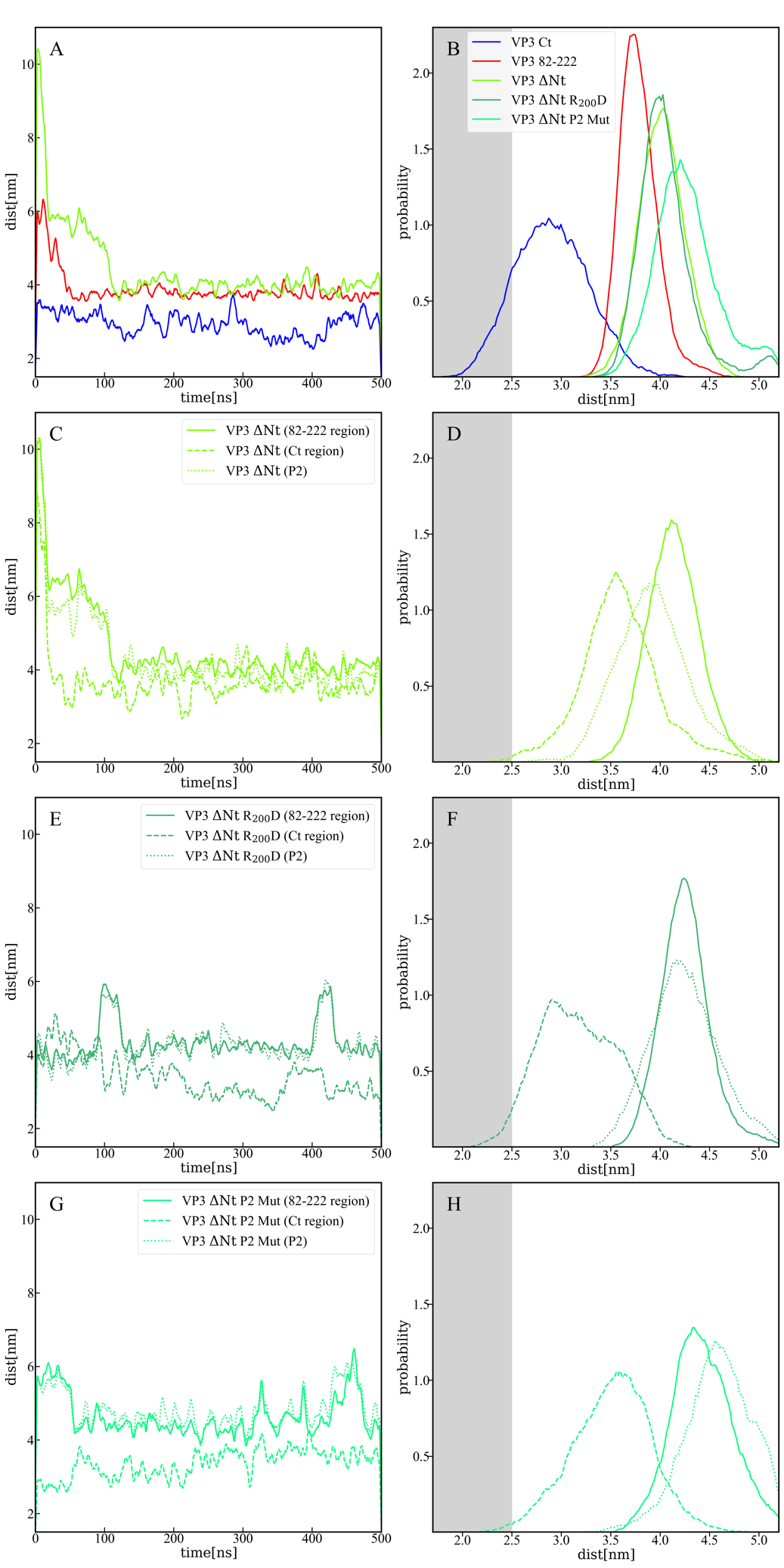

**Fig. S10. Distance between membrane and protein constructs** **computed by MD**.

**(A)** Time evolution of the distance (perpendicular to the membrane surface) between the center of mass of VP3 Ct (blue solid line), VP3 82-222 (red solid line), and VP3 ΔNt (green solid line) and the center of mass of the membrane.

**(B)** Distribution of distances computed from the time-traces of panel A in the time interval [200-500] ns; and also from simulations of two mutants of VP3 ΔNt: the single mutant R_200_D (VP3 ΔNt R_200_D) and the one with the four residues of the P2 region mutated K_157_D, R_159_D, H198D and R_200_D (VP3 ΔNt P2 Mut). The VP3 Ct construct gets closer to the membrane than any other construct or protein domain and, more importantly, VP3 ΔNt adsorbs with the Ct region facing the negatively charged membrane. The single mutant exhibits a binding similar to wild type VP3 ΔNt protein, while the one with the whole P2 mutated is located further away from the negative surface. Results obtained by coarse-grained Molecular Dynamics simulation of VP3 Ct, VP3 82-222, and VP3 ΔNt, in front of a DOPE, DOPC, PI3P membrane (molar ratios of 64:31:5).

**(C, E, G)** Time evolution of the distance between the center of mass of different protein regions and the center of mass of the membrane for VP3 ΔNt and the two mutants. The continuous line corresponds to the center of mass of the region 82-222, the dashed lines, to the Ct region, and the dotted lines, to the P2 region.

**(D, F, H)** Distribution of distances computed from the time-traces of panels C, E and G in the time interval [200- 500] ns.

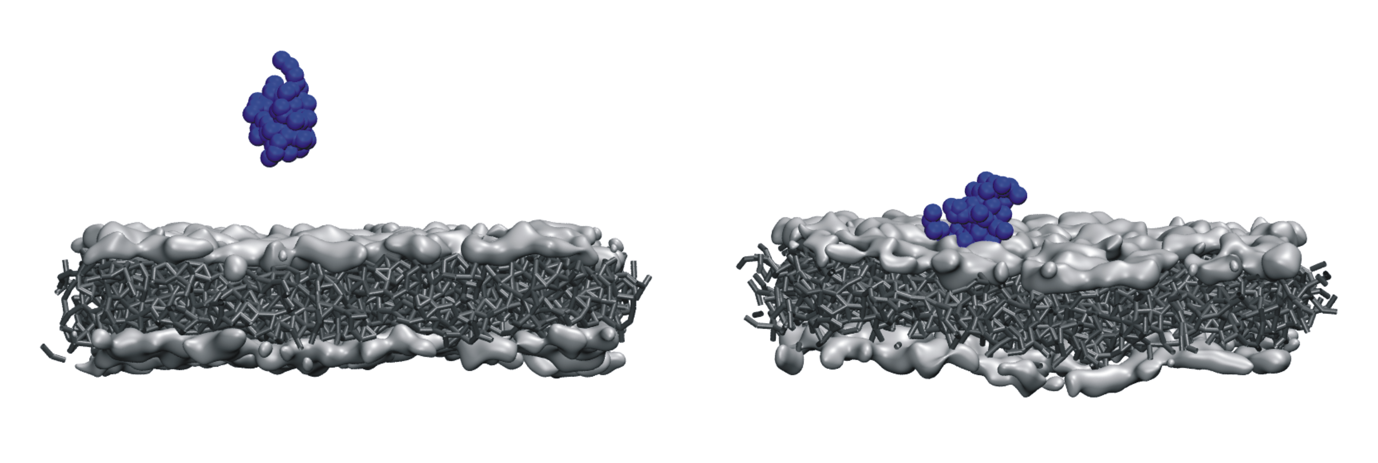

**Fig. S11. Initial and final configurations of VP3 Ct on a lipid membrane.**

Initial (left panel) and final (right panel) configurations from a 500 ns coarse-grained Molecular Dynamics simulation of VP3 Ct in front of a DOPE, DOPC, PI3P membrane (molar ratios of 64:31:5).

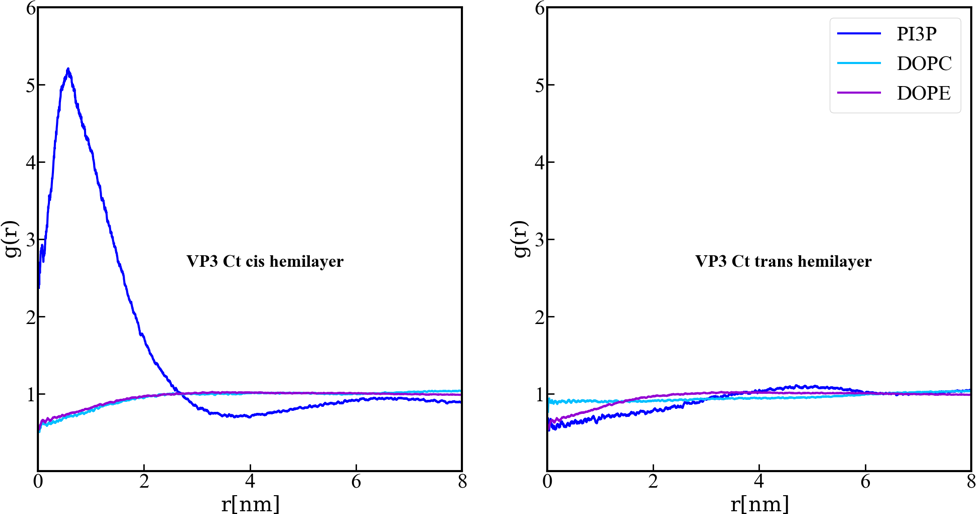

**Fig. S12. Lipids accumulation/depletion around VP3 Ct.**

In plane (X-Y) radial distribution functions, g(r). These functions were computed during the 200-500 ns time interval of the Molecular Dynamics simulations described in the text and in the previous supporting figures. The panels display the g(r) between the lipids in the cis- (left) and trans- (right) membrane hemilayers, and the center of mass of the VP3 Ct construct. Once again, this positively charged construct attracts the PI3P molecules from the cis-hemilayer to its proximity, as evident from the values of g(r)>1 for r<2 nm. Moreover, for VP3 Ct this effect is strong enough to deplete the PI3P molecules in the trans-hemilayer due to inter-hemilayer PI3P-PI3P electrostatic repulsions.

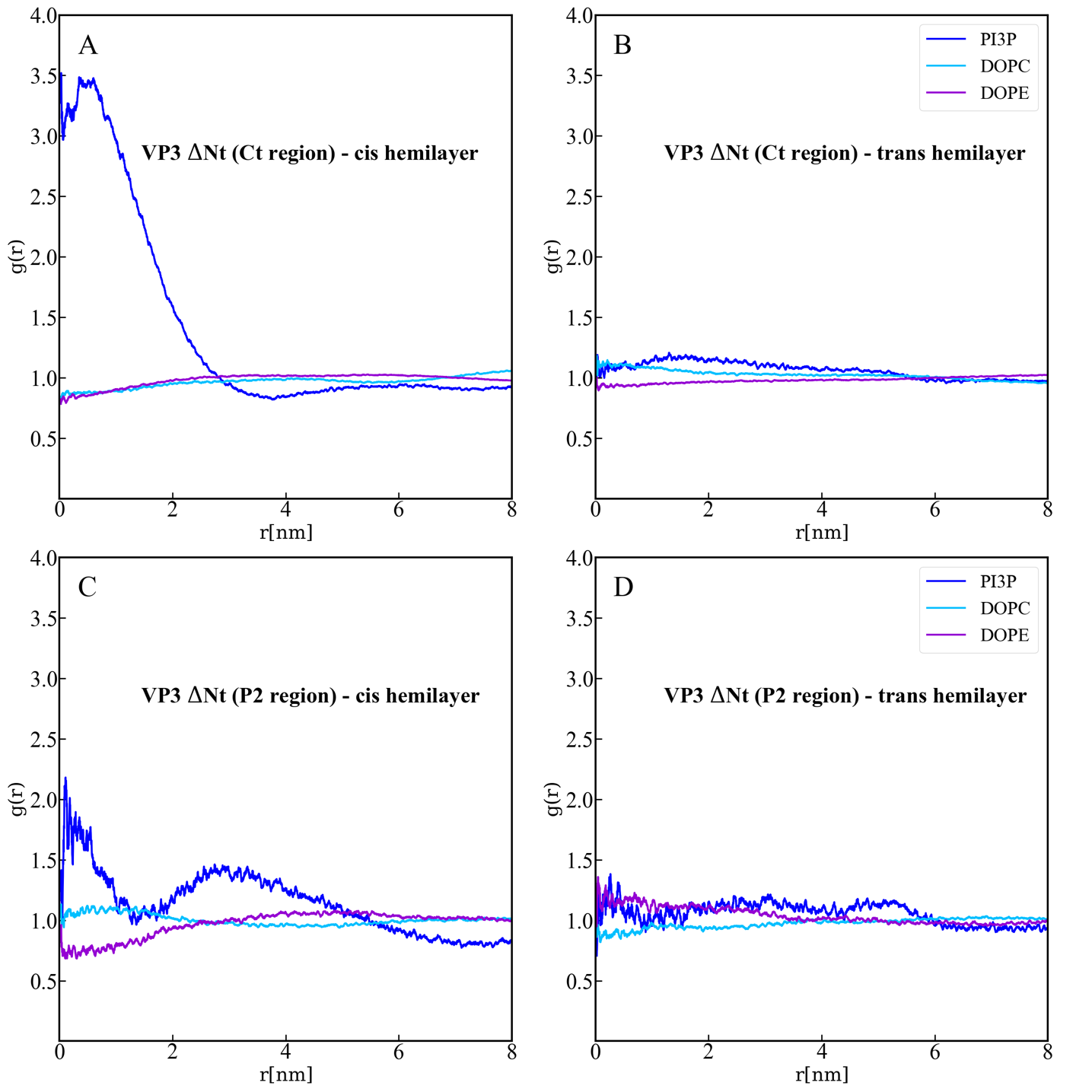

**Fig. S13. Lipids accumulation/depletion around the Ct fragment of VP3 ΔNt.**

In plane (X-Y) radial distribution functions, g(r). These functions were computed during the 200-500 ns time interval of the Molecular Dynamics simulations described in the text and in the previous supporting figures. The upper panels display the g(r) between the lipids in the cis- (left) and trans- (right) membrane hemilayers, and the center of the Ct domain of VP3 ΔNt. As observed, the Ct domain attracts PI3P to its proximity, within a radius of approximately 4 nm where g(r)>1 (blue line), in the hemilayer that is in contact with the protein (cis-). The lower panels depict the radial distribution functions between the lipids in the cis- (left) and trans- (right) hemilayers, and the P2 domain of VP3 ΔNt. Once again, P2 attracts PI3P to its proximity in the cis-hemilayer, within a radius of about 5 nm.

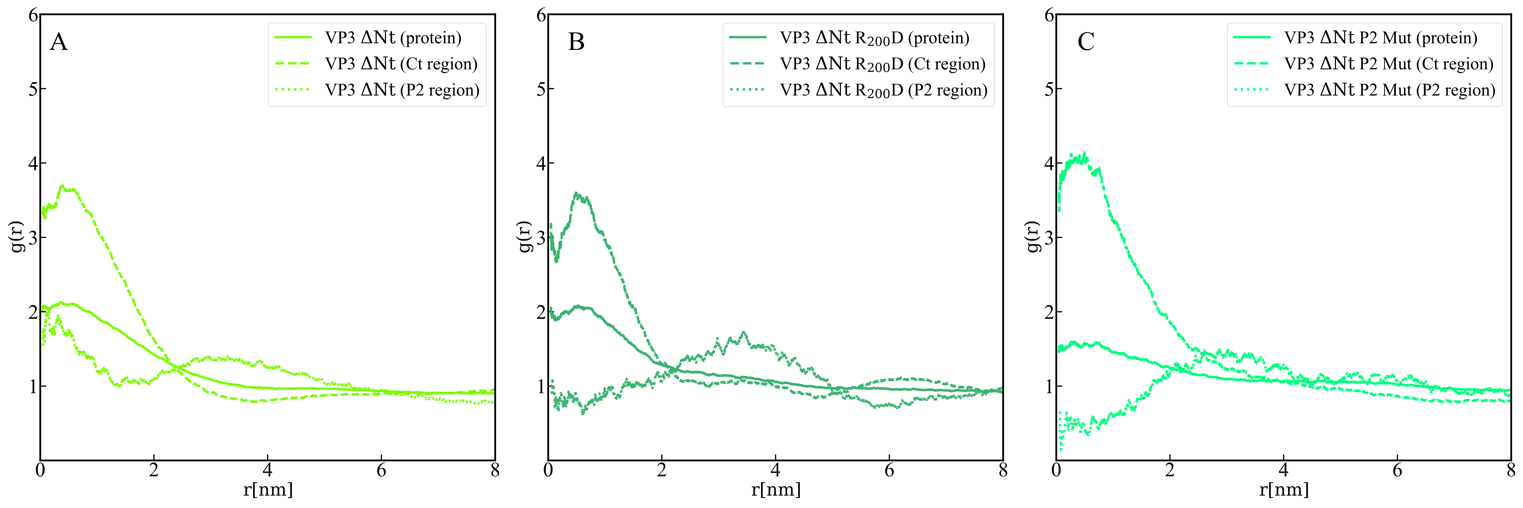

**Fig. S14. Lipids accumulation/depletion around VP3 ΔNt and its mutants.**

In plane (X-Y) radial distribution functions, g(r) computed in the 200-500 ns time interval of the Molecular Dynamics simulations of the VP3 ΔNt protein (A) and two mutants: VP3 ΔNt R_200_D (B) and VP3 ΔNt P2 Mut (C). The panels display the g(r) between PI3P lipids in the *cis* membrane hemilayer, and the center of mass of the whole protein (continuous line), the Ct region (dashed line) and the P2 region (dotted line). The concentration effects on PI3P, induced by VP3 ΔNt, is exerted by the positively charged Ct and the P2 regions. When the P2 region is mutated, the P2 contribution to PI3P recruitment is diminished gradually from the single mutant to the wholly mutated P2.

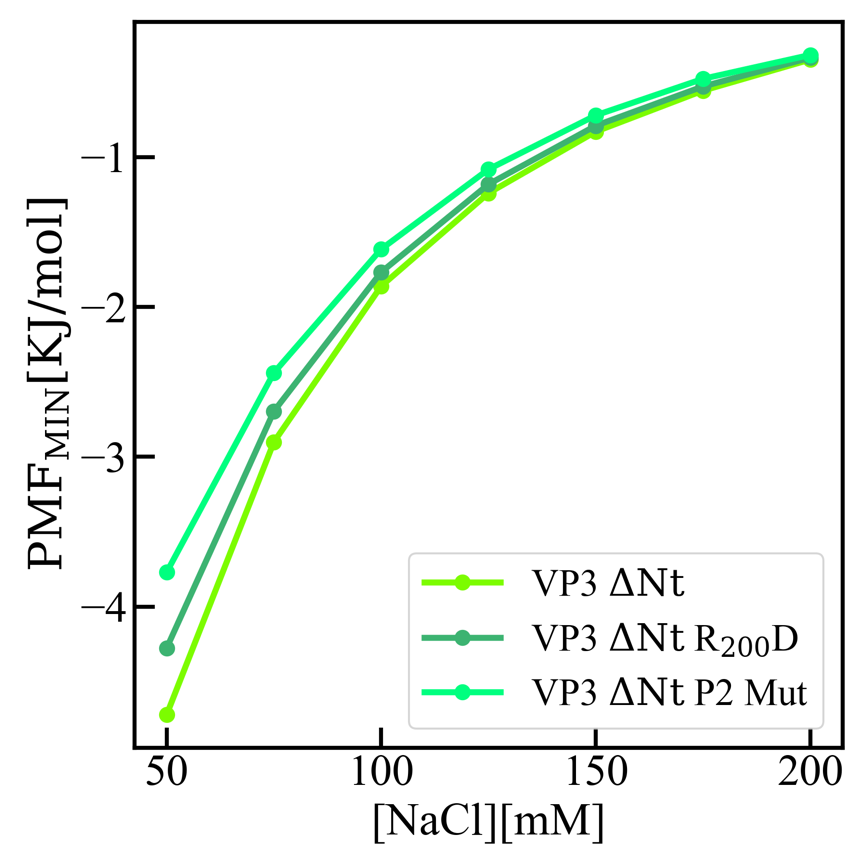

**Fig. S15. Adsorption free energy of VP3 ΔNt and VP3 ΔNt mutants on PI3P(+) model membranes.** Adsorption free energy (ΔG_ad_), computed from the minimum of PMF(z), versus concentration of NaCl, for VP3 ΔNt and the mutants VP3 ΔNt R_200_D and VP3 ΔNt P2 Mut. In all cases, the solution pH is 8, and the concentration of protein is 1 µM. The membrane surface contains 5% titratable groups, representing PI3P. Each group has three acidic moieties, one with a pKa of 2.5 and the others with 6.5. At 150 mM NaCl and pH 8, more than 90% of the acidic groups are deprotonated. As the number of positively charged residues in VP3 is systematically reduced from VP3 ΔNt to VP3 ΔNt P2 Mut, the binding of the protein to the membrane becomes weaker. The effect is more pronounced at lower salt concentrations, which highlights the weight of electrostatic forces on the adsorption of VP3 on negatively charged membranes.

**Table S1. Primers**

| **Construction** | **Name of primer pairs** | **Sequences*** | **Position, mutation introduced or fragment** |
| --- | --- | --- | --- |
| pcDNA VP3 FL K157D | VP3.157 | 5’ ATGCAGAG***G***A***C***AGCCGGTTGGCATC 3’ (sense)  5’ AACCGGCT***G***T***C***CTCTGCATGCAC 3’ (antisense) | 157, K to D |
| pcDNA VP3 FL R159D | VP3.159 | 5’ GCATGCAGAGAAGAGC***GAC***TTGGCATCAG 3’ (sense)  5’ ***GTC***GCTCTTCTCTGCATGCACGTAGTCTAG 3’ (antisense) | 159, R to D |
| pcDNA VP3 FL H198D | VP3.198 | 5’ GTCTATGAAATCAAC***G***ATGGACGTGGC 3’ (sense)  5’ ***C***GTTGATTTCATAGACTTTGGCAACTTC 3’ (antisense) | 198, H to D |
| pcDNA VP3 FL R200D | VP3.200 | 5’ GAAATCAACCATGGA***GA***TGGCCCAAAC 3’ (sense)  5’ ***TC***TCCATGGTTGATTTCATAGACTTTGG 3’ (antisense) | 200, R to D |
| pFastBacHTb-his-VP3 FL | VP3.Fw  VP3.Rv | 5’ GGATCCGCTGCATCAGAGTTCAAAGAGAC 3’  5’ GAATTCTCACTCAAGGTCCTCATCAGAG 3’ | VP3 FL |
| pFastBacHTb-his-VP3 ∆223-257 | VP3.Fw  VP3.83-222.Rv | 5’ GGATCCGCTGCATCAGAGTTCAAAGAGAC 3’  5’ GAATTCTCAATTGCGATGCTTCATCTC 3 3’ | VP3 ∆223-257 |
| TrxA.His.Ts.2xFYVE | 2xFYVE | 5’ ACCGACGACGACGACAAGGAAAGTGATGCCATGTT 3’ (sense)  5’ GTGGTGGTGGTGCTCGAGGCCCGCGGTACCGTCGA 3’ (antisense) | 2xFYVE |
| TrxA.His.Ts.VP3FL | VP3.FL.Fw  VP3.FL.Rv | 5’ CTGGTGCCACGCGGTTCTGCATCAGAGTTCAAAGA 3’  5’ GCCCGCGGTACCGTCGACTTACTCAAGGTCCTCAT 3’ | VP3 FL |
| TrxA.His.Ts.VP3∆223-257 | VP3.FL.Fw  VP3.∆223257.Rv | 5’ CTGGTGCCACGCGGTTCTGCATCAGAGTTCAAAGA 3’  5’ GCCCGCGGTACCGTCGACTCAATTGCGATGCTTCA3’ | VP3 ∆223-257 |

* For the first four point mutants, the nucleotides that allow to introduce amino acid changes in the VP3 protein are indicated in italic bold letters. The underlined nucleotides indicate restriction sites.
